## Supplementary material for "SLC35G3 is a UDP-N-acetylglucosamine transporter for sperm glycoprotein formation and underpins male fertility in mice": SI

**Affiliation:**




**This PDF file includes:**

Figs. S1 to S12

**Other Supplementary Materials for this manuscript include the following:**

Tables S1 to S3

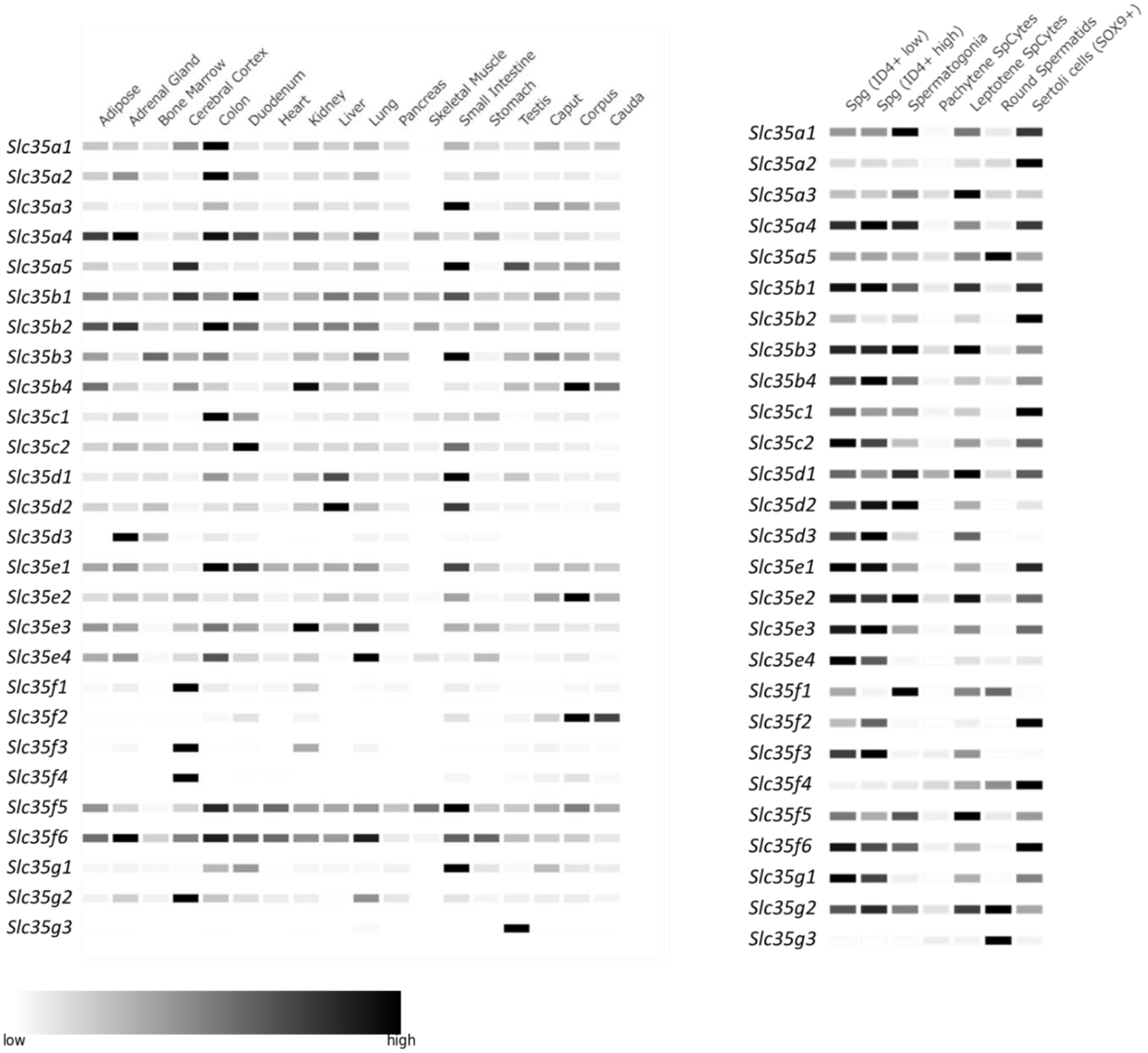


**Fig. S1. *Slc35g3* is the sole Slc35 family member that is testis-specific, with highest mRNA levels in round spermatids.**

Levels of mRNA for Slc35 family genes in organs and testis ages was created using Mammalian Reproductive Genetics Database V2.

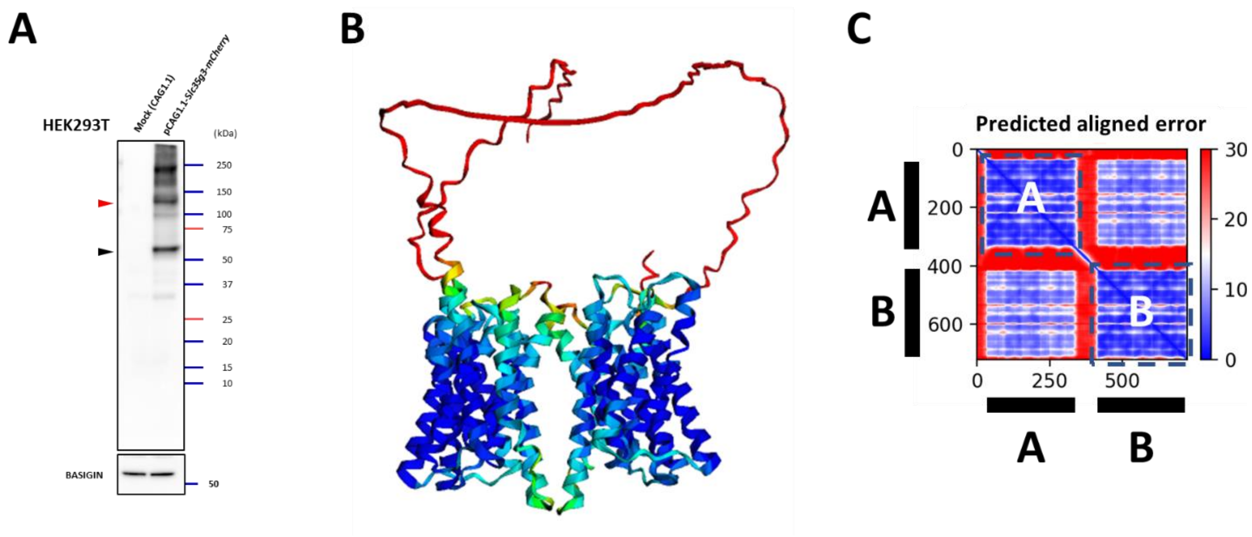


**Fig. S2. SLC35G3 has the potential to form a homodimer**

**(A)** Transfection of pCAG1.1-Slc35g3-mCherry into HEK293T cells followed by Western blot analysis using an mCherry antibody revealed a signal at the predicted size of 62.2 kDa (black arrowhead). Simultaneously, a signal was also detected at approximately twice the size (red arrowhead). **(B)** The predicted SLC35G3 structure using AlphaFold2 shows a potential homodimeric arrangement. **(C)** The predicted aligned error map from AlphaFold2 indicates that the aligned errors for both SLC35G3 proteins, denoted as A and B, are within 15Å. The predictions for A-B and B-A interactions are also below 15Å, suggesting a plausible structural model for SLC35G3 homodimer formation.

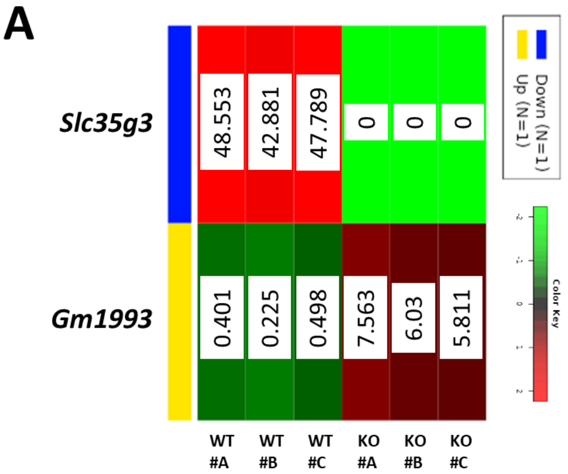


B

| Symbol | log2 FC | Adj.Pval | Chr | feature |
|---|---|---|---|---|
| ***Slc35g3*** | -5.56 | 6.55e-06 | 11q | Analyzed in this study |
| ***Gm1993*** | 2.44 | 1.56e-02 | Xq | Gm1993 is related to Sycp3. Sycp3 KO: male infertility; Meiosis in males blocks at early prophase with lack of synaptonemal complexes. |

**Fig. S3. Differentially expressed genes (DEG) analysis using testis RNA-seq data**
**(A)** Genes called by RNA-seq using wild-type (WT; n=3) and knockout (KO; n=3) testis followed by DEG analysis. Blue indicates a gene that is decreased in KO (*Slc35g3*), and yellow indicates a gene that is increased in KO (*Gm1993*) compared to WT. Numbers indicate FPKM. **(B)** Log2 Fold Change, *P* value, gene locus, and feature of each DEG are shown.

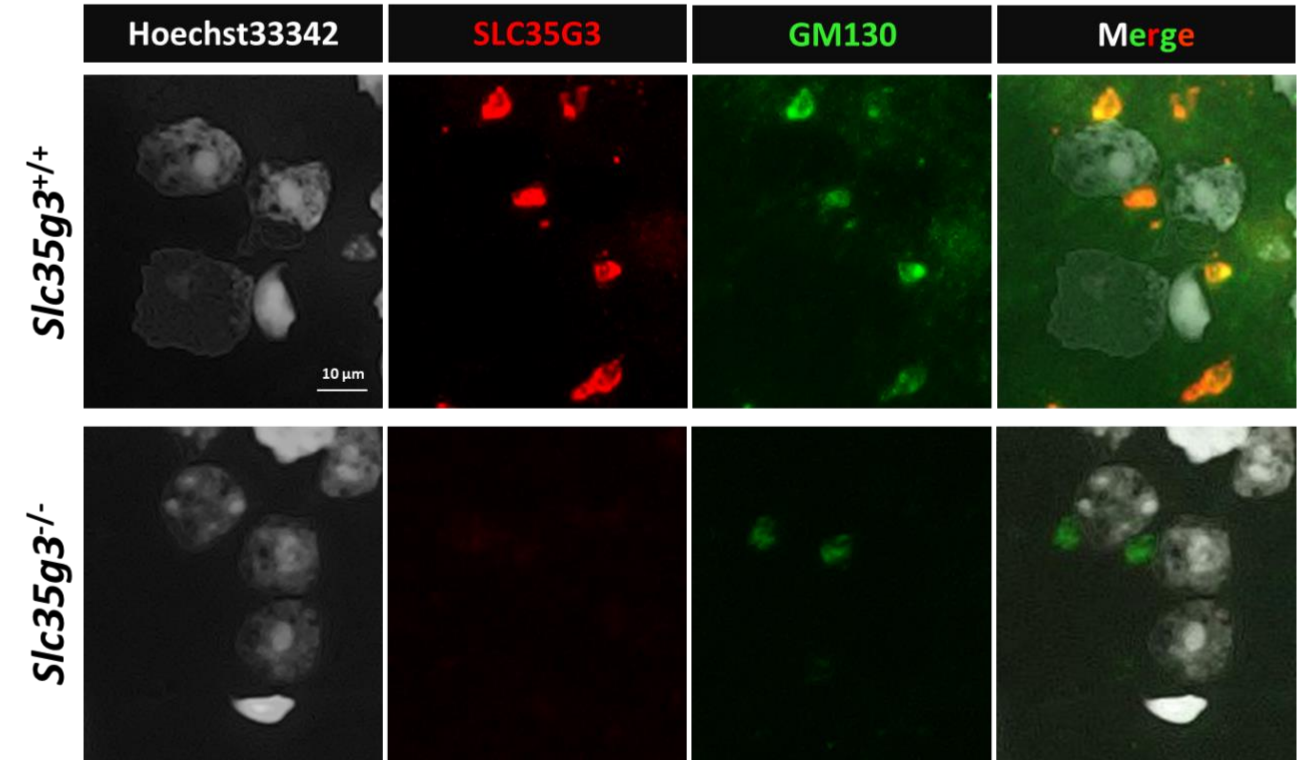


**Fig. S4. SLC35G3 is localized in the Golgi apparatus, and the signal disappears in the knockout.**

The top row displays immunostaining images of *Slc35g3*$^{+/+}$ testicular cells, while the bottom row displays immunostaining images of *Slc35g3*$^{-/-}$ testicular cells. Each image shows the cell nucleus obtained by crushing the seminiferous tubule (Hoechst33342), SLC35G3, GM130, and a merged image. The upper panels are the same as **Fig. 1E**.

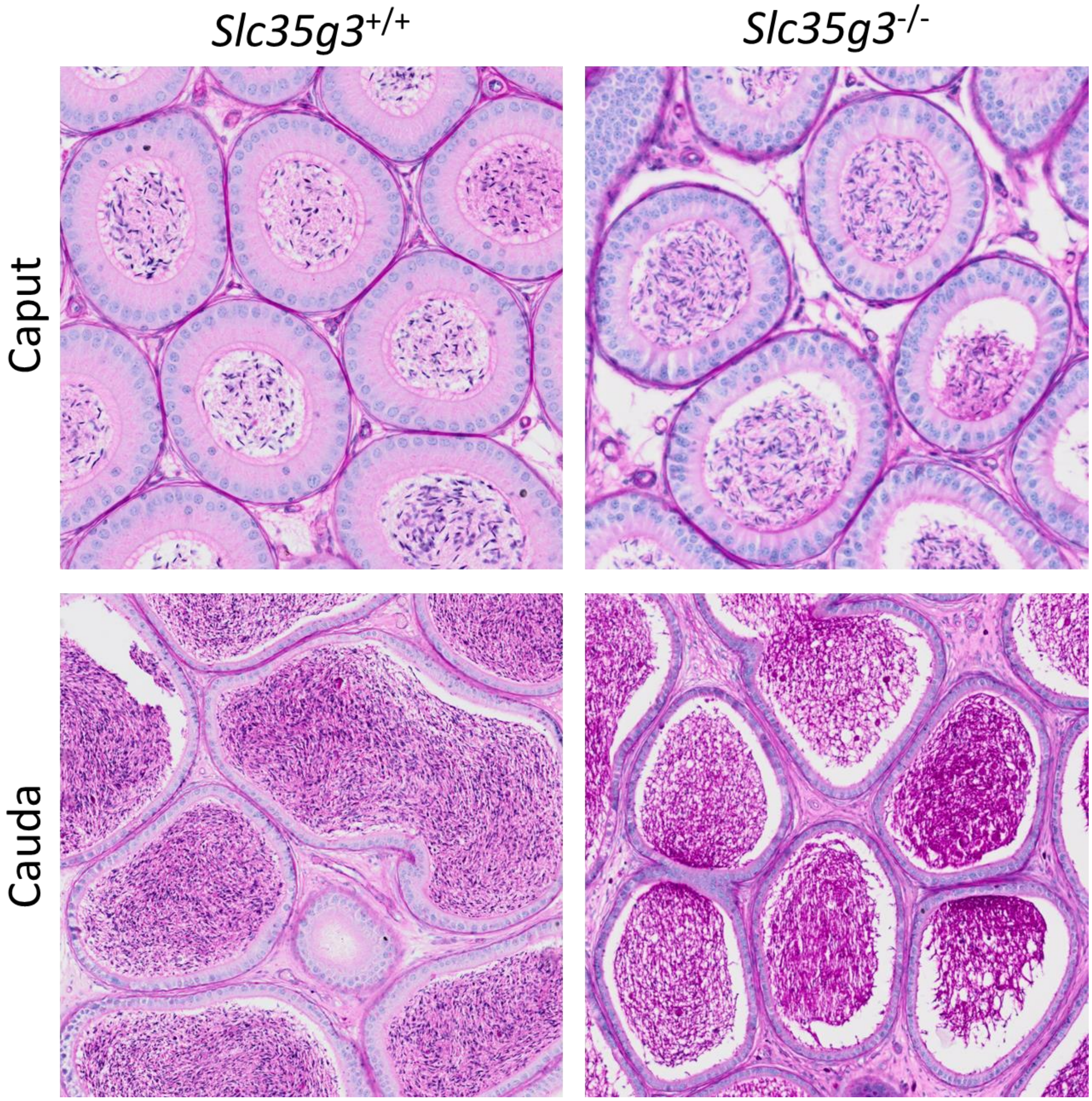


**Fig. S5. Epididymis sections from *Slc35g3*$^{+/+}$ and *Slc35g3*$^{-/-}$ mice**

Caput and cauda epididymis sections from *Slc35g3*$^{+/+}$ and *Slc35g3*$^{-/-}$ mice are comparable.

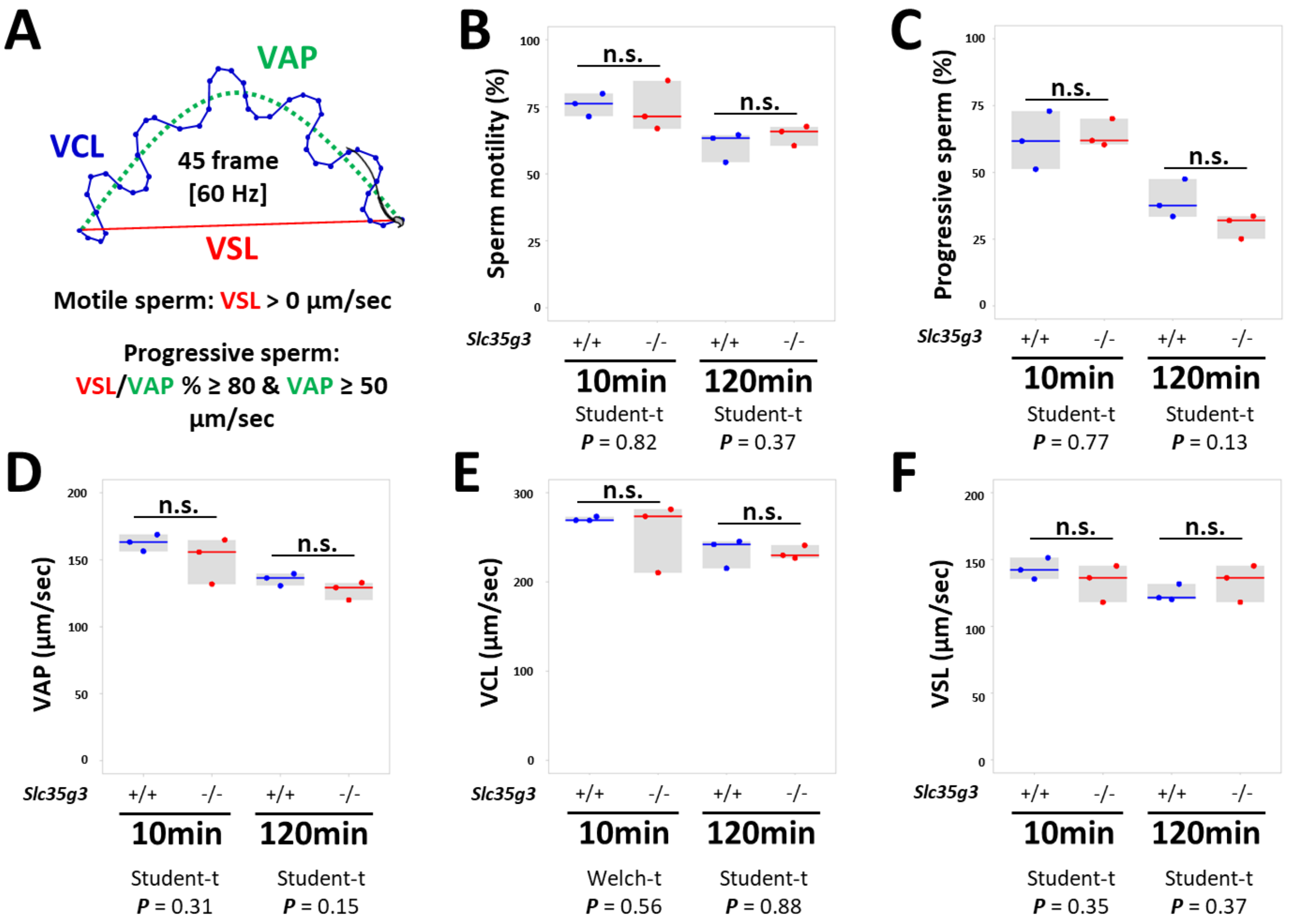


**Fig. S6. No significant differences in CASA parameters of sperm from *Slc35g3*$^{-/-}$ mice compared to sperm from *Slc35g3*$^{+/+}$ mice.**

**(A)** Definition of parameters representing sperm motility; Velocity of average path (VAP) is indicated in green, Velocity of curved path (VCL) is indicated in blue, and Velocity of straight path (VSL) is indicated in red. Motile sperm is defined as having a VSL greater than 0 µm/sec, while progressive sperm is defined as having VSL/VAP % greater than or equal to 80 and VAP greater than or equal to 50 µm/sec. **(B)** Motility of sperm from wild-type (+/+) and knockout (-/-) mice showed no significant (n.s.) differences at 10 and 120 minutes after suspension in the medium. **(C)** Percentage of progressive sperm from *Slc35g3*$^{+/+}$ and *Slc35g3*$^{-/-}$ mice showed no significant differences at 10 and 120 minutes after suspension in the medium. **(D)** VAP of wild-type and knockout sperm showed no significant differences at 10 and 120 minutes after suspension in the medium. **(E)** VCL of wild-type and knockout sperm showed no significant differences at 10 and 120 minutes after suspension in the medium. **(F)** VSL of wild-type and knockout sperm shows no significant differences at 10 and 120 minutes after suspension in the medium.

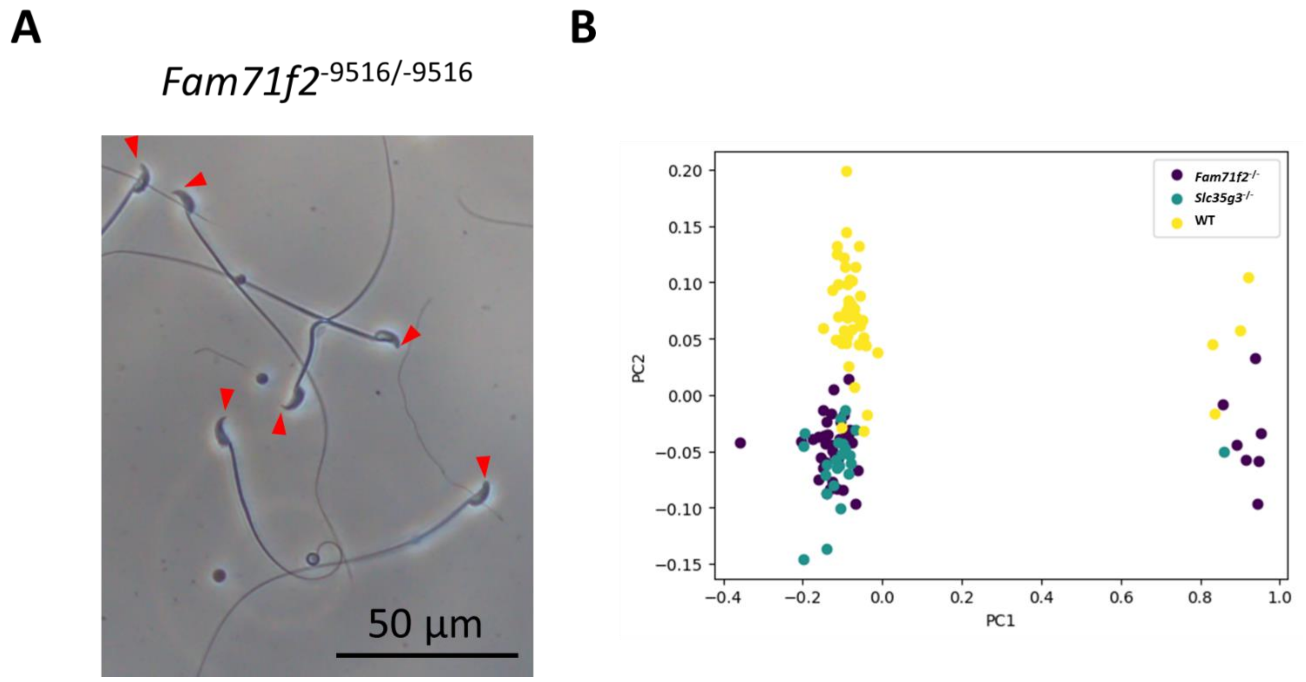


**Fig. S7. Sperm from *Slc35g3*$^{-/-}$ mice exhibit morphology similar to sperm from *Fam71f2*$^{-/-}$ mice**

**(A)** Appearance of *Fam71f2* KO sperm. Red arrowheads indicate the tips of the spermatozoa. **(B)** Comparison among sperm from wildtype, *Fam71f2*$^{-/-}$, and *Slc35g3*$^{-/-}$ mice by elliptic Fourier analysis.

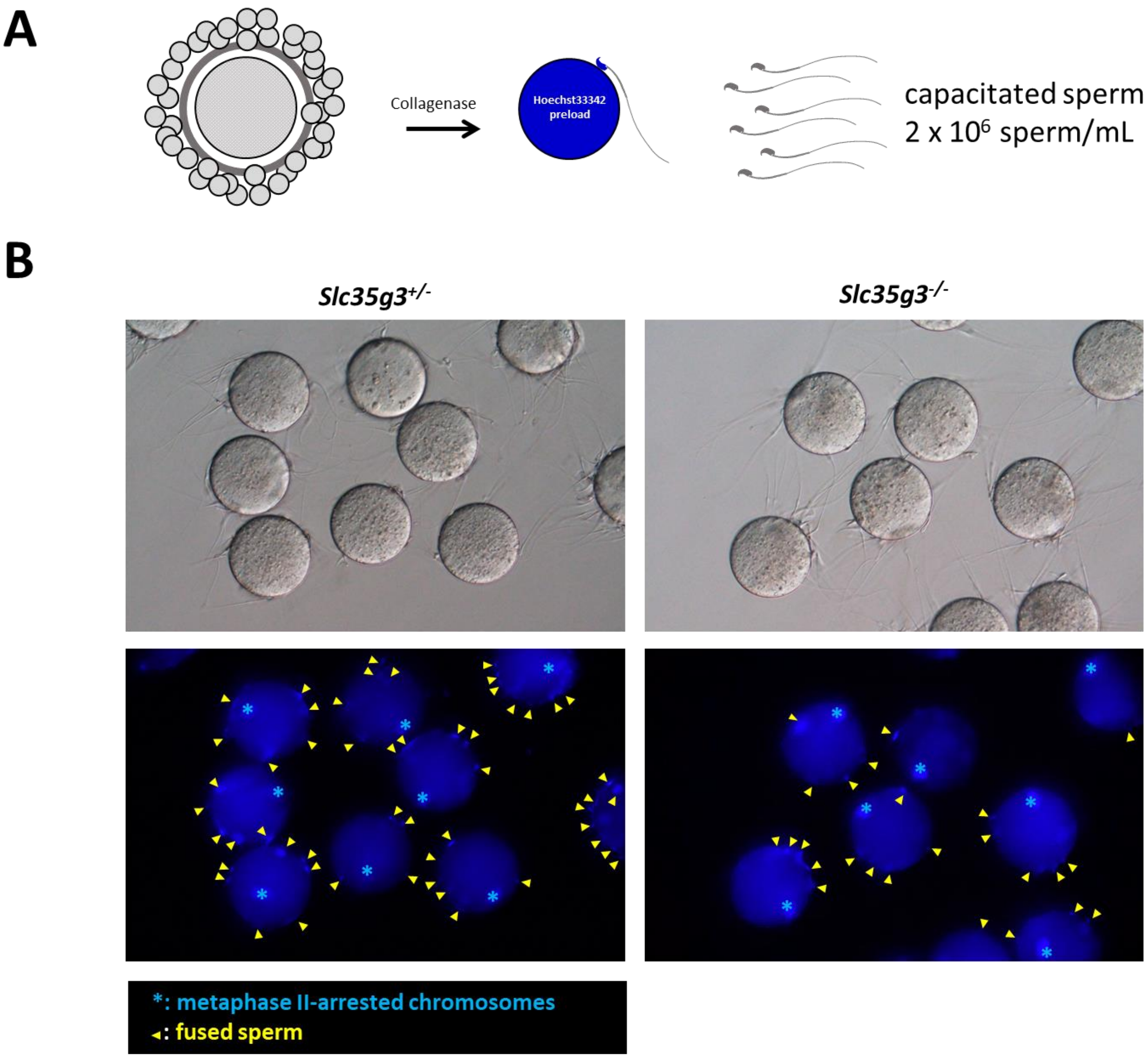


**Fig. S8. Oolema fusion observed under high sperm concentration conditions.**
**(A)** Schematic diagram of fusion test using 2 x $10^6$ sperm/mL. **(B)** Bright Field and Hoechst stained images. The yellow arrowheads show fused spermatozoa and sky-blue asterisks shows metaphase II-arrested chromosomes.

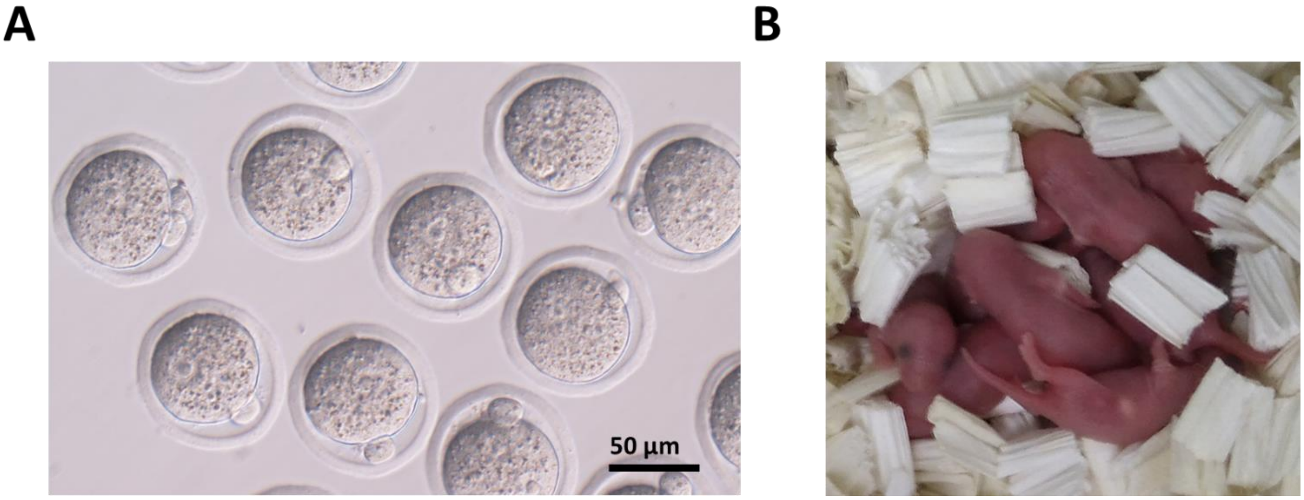


**Fig. S9. Pups were produced from spermatozoa from *Slc35g3*$^{-/-}$**
**(A)** Two-pronuclear embryos were obtained by IVF using 2 x $10^6$ sperm. **(B)** Offspring obtained by embryo transfer.

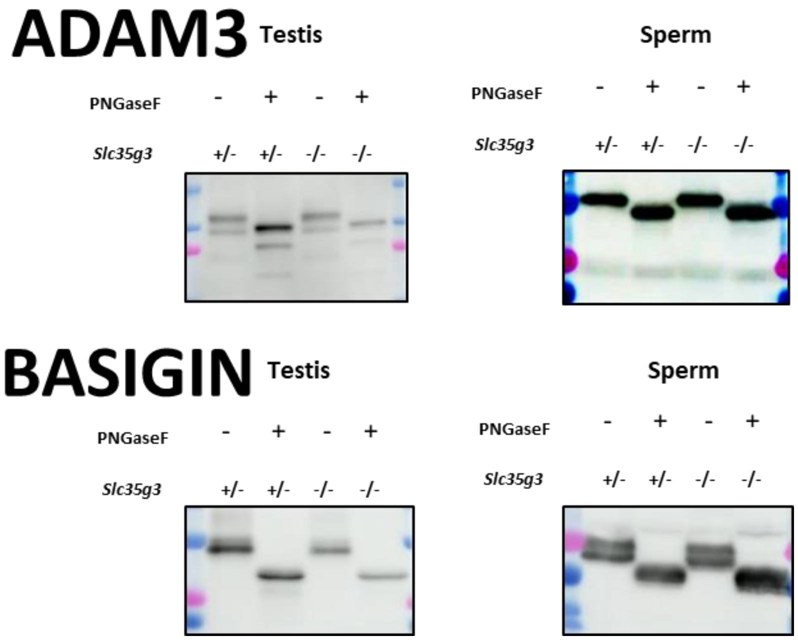


**Fig. S10 ADAM3 patterns were comparable between Slc35g3$^{+/-}$ and Slc35g3$^{-/-}$ after PNGaseF treatment**

Western blot analysis of PNGaseF treated or non-treated ADAM3 from testis and sperm of *Slc35g3*$^{+/-}$ and *Slc35g3*$^{-/-}$ mice. BASIGIN was analyzed as a loading control.

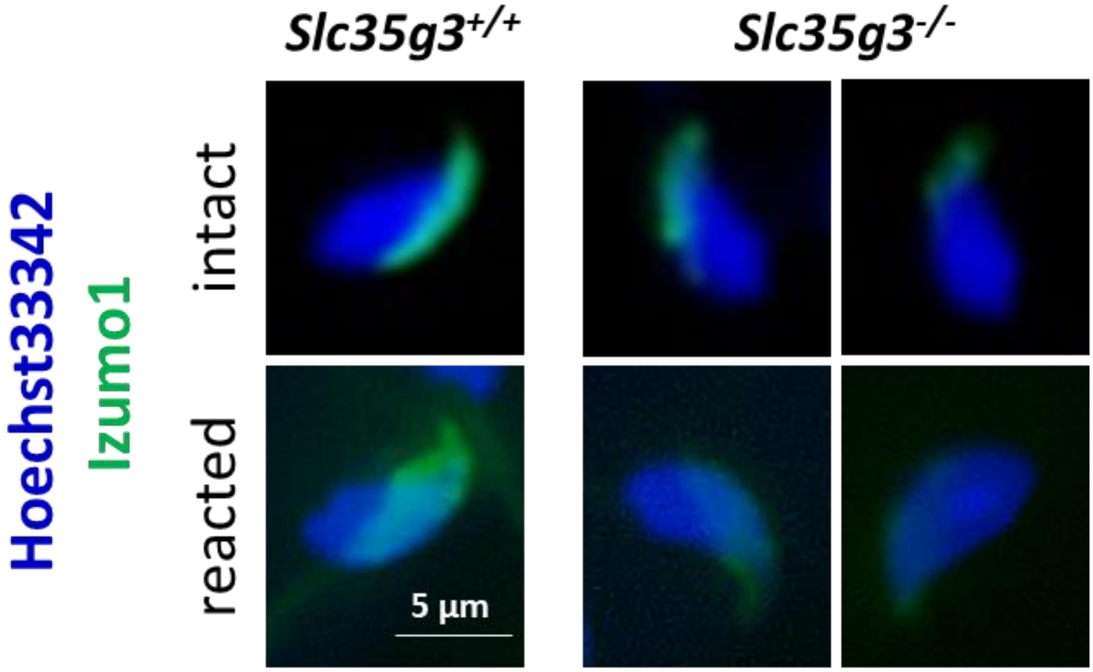


**Fig. S11. Distribution of IZUMO1 is expanded in sperm from Slc35g3$^{-/-}$ mice after the acrosome reaction**

Acrosome-intact sperm (upper row) and acrosome-reacted sperm (lower row) from *Slc35g3*$^{+/+}$ mice (left column) and *Slc35g3*$^{-/-}$ mice (right column).

.

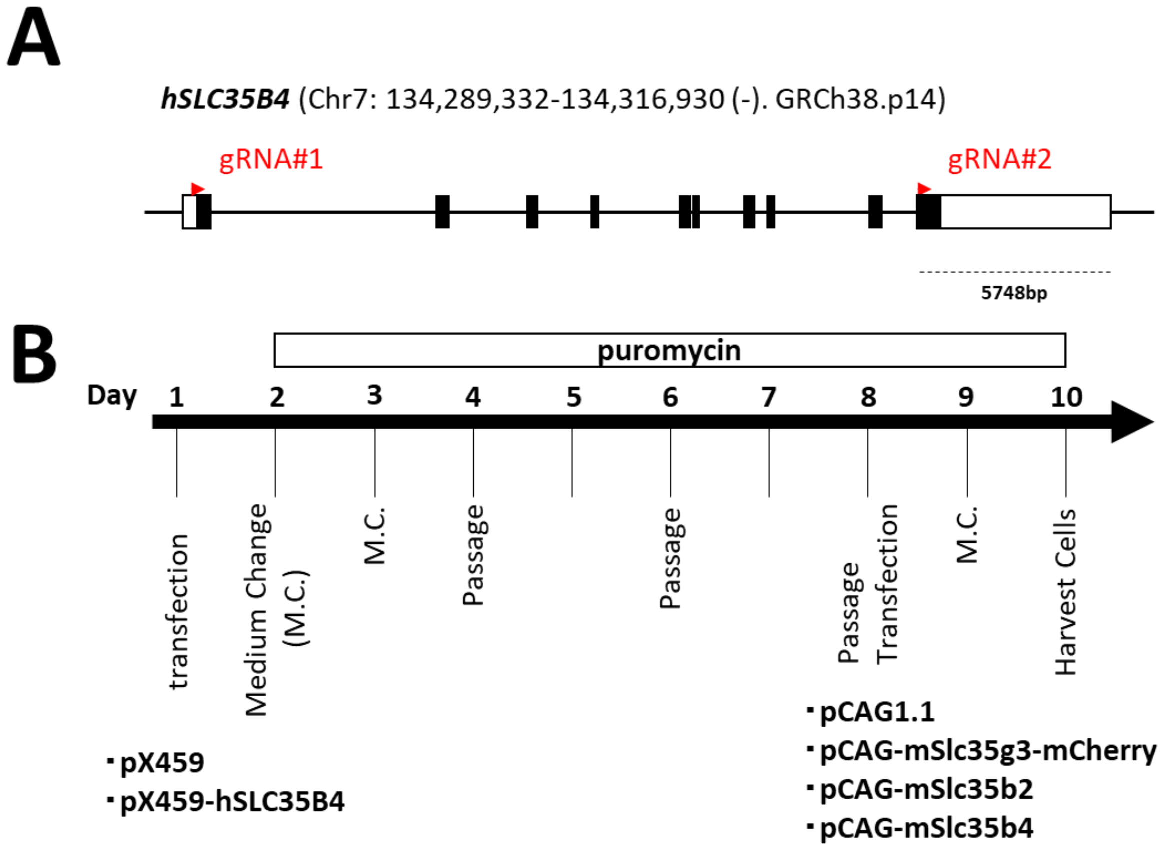


**Fig. S12. Design of gRNAs and timeline of the experiment**

**(A)** The gRNAs were designed to remove almost the entire coding region of the exon.
**(B)** After the introduction of pX459, selection was performed using puromycin as shown.

Table. S1. Mass spectrometry data using sperm lysates from *Slc35g3*$^{+/+}$ and *Slc35g3*$^{-/-}$ mice.

Table. S2. Primer and gRNA sequences used in the present studies.

Table. S3. Antibodies used in the present studies.
